# Heme concentration-dependently modulates the production of specific antibodies in murine

**DOI:** 10.1101/071316

**Authors:** Guofu Li, Haiyan Xue, Zeng Fan, Yun Bai

**Affiliations:** School of Life Sciences, Sun Yat-Sen University, Guangzhou, China, 510275

**Keywords:** Heme, Specific antibody, Inflammation, Innate immunity, Adaptive immunity

## Abstract

Free heme is an endogenous danger signal to provoke innate immunity. Active innate immunity is a precondition of an effective adaptive immune response. However, heme catabolites, CO, biliverdin and bilirubin trigger immunosuppression. Furthermore, free heme induces the expression of heme oxygenase-1 to reinforce the production of CO, biliverdin and bilirubin. As such, free heme can drive two antagonistic mechanisms to affect adaptive immunity. What is the outcome of animal immune response to an antigen in the presence of free heme? The question remains to be explored. Here we report the immunization results by intraperitoneal injection of the formulations containing BSA and heme. When the used heme concentrations were about less than 1 µM, the production of anti-BSA IgG and IgM was unaffected; when the used heme concentrations were about more than 1 µM but less than 5 µM, the production of anti-BSA IgG and IgM was enhanced; when the used heme concentrations were about more than 5 µM, the production of anti-BSA IgG and IgM was suppressed. The results demonstrate that heme can modulate adaptive immunity (at least humoral immunity) by the mode of double concentration-thresholds. If heme concentrations are below the first threshold, there is no effect on adaptive immunity; if between the first and second thresholds, there is promotive effect; if over the second threshold, there is inhibitive effect. A hypothesis is also presented here to explain the mode.

**Abbreviations**
CFA: Complete Freund’s adjuvant
DAMP: damage associated molecular pattern
PAMP: pathogen-associated molecular pattern
HO: heme oxygenase
HEcL: human erythrocyte lipid
REcL: rat erythrocyte lipid
LREcL: hemolytic rat erythrocyte lipid
LPS: Lipopolysaccharide
PC: phosphatidylcholine
ROS: reactive oxygen species
TLR: Toll-like receptor

## 1. Introduction

Heme refers to several iron porphyrins in organisms, such as heme a, b and c. Heme b is the basic biosynthetic type and others are its subtle derivatives. Free heme can do harm to cells through directly wedging into biological membrances and modifying proteins, DNAs and lipids by producing reactive oxygen species (ROS) [1]. So the most heme is in the protein-bound form and the concentration of free heme is strictly controlled under physiological condition[2]. Red blood cells and muscle cells contain the most heme of an animal and tend to raise the level of free heme due to the cell renewal or injury. To constrain the level of free heme, besides balancing its synthesis and degradation, animals evolutionarily develop a buffer system composed of haptoglobin, hemopexin and some other serum proteins[2]. Haptoglobin, a serum protein primarily produced from liver, binds hemoglobin or myoglobin to form a high-affinity complex that will then be removed by the reticuloendothelial system. Hemopexin, another serum protein synthesized by liver, has the highest affinity for heme, binds and delivers heme to the liver for further catabolism. ther serum proteins such as serum albumin can do the same works as hemopexin.

Heme plays versatile roles in the process from cell division, differentiation, apoptosis and necrosis to individual development, growth and diseases. The versatile roles rely on the heme-controlled biomolecule network covering signal transduction, gene expression and metabolism. The elements for heme to weave the network are hemoproteins, heme-responsive proteins and heme-dependent ROS[1]. First, the major form of heme in organisms is as the prosthetic group of numerous hemoproteins to carry out diverse functions[3, 4], including electron transporters such as cytochromes in electron transport chains[5], gas carriers such as hemoglobin for O_2_, gas sensors such as CooA for CO, FixL for O_2_ and soluble guanylate cyclase for NO[6], and oxidation/reduction enzymes such as cytochrome p450[7]. Second, free heme is a regulator of numerous heme-responsive proteins[4, 8]. Heme-responsive proteins can be protein kinases[9, 10], transcription factors[11, 12], ion channels[13, 14], microRNA processing factors[15, 16] and so on. In the third place, free heme produces ROS through NADPH oxidases[17] and Fenton reaction[18]. ROS regulates a diverse array of signaling pathways by controlling the thiol/disulfide redox states of proteins[19].

Heme is also intensively involved in immune system but acts paradoxically[20]. On the one hand, free heme activates toll-like receptor 4 (TLR4) dependent[21-23] and ROS dependent signaling pathways[24-26]. The activation of these signaling pathways promotes leukocyte maturation/migration[27-29] and anti-apoptosis[30, 31], pro-inflammatory cytokine secretion[32], adhesion molecule expression[33, 34] and ROS production[28, 35], all of which construct a vigorous innate immune response or inflammation[36, 37]. On the other hand, there have been many reports about the negative effect of heme on innate immunity or inflammation, such as promoting cell apoptosis[38, 39] and anti-inflammatory cytokine secretion[40]. The suppressive mechanism depends on heme oxygenases (HO), including inducible HO-1 and constitutive HO-2. HO is the key rate-limiting enzyme to convert heme into Fe^2+^, CO and biliverdin which is subsequently transformed to bilirubin by biliverdin reductase[20]. CO, biliverdin and bilirubin trigger anti-inflammatory/immunosuppressive signaling pathways [41-43]. Furthermore, free heme induce the expression of HO-1 to reinforce the anti-inflammatory/immunosuppressive signaling pathways [44]. HO-1 can be induced by many other stressors and becomes a hot target for various anti-inflammatory or immunosuppressive therapies[45].

Innate immunity controls adaptive immunity[46, 47], so free heme has the potential to regulate adaptive immunity. However, what is the final effect of the self-contradictory free heme on animal immune response to an antigen, promotion, inhibition or no influence? The question remains to be explored. We here report the effect of free heme on the production of anti-BSA antibodies in rats and mice.

## 2. Materials and Methods

### 2.1. Animals

Sprague Dawley (SD) rats and BALB/c mice, including males and females, were obtained from the animal center of Sun Yat-sen University. All the rats and mice were housed in individual cages with free access to sterile water and irradiated food in a specific pathogen-free facility. Animal experiments were conducted in accordance with the institutional guidelines of Sun Yat-sen University.

### 2.2. Chemicals and reagents

Limulus amebocyte lysate (LAL) test reagents were purchased from Associates of CAPE COD. Bovine serum albumin (BSA), egg phosphatidylcholine (PC), Complete Freund’s adjuvant (CFA) and heme (hemin, HPLC grade purity > 98.0%) were purchased from Sigma-Aldrich. LAL test showed that the lipopolysaccharide (LPS) activity of 250 µM heme solution was less than 0.01 EU/ml. HRP labeled ant-rat IgG (H/L), ant-rat IgM (μ), ant-mouse IgG (H/L), anti-mouse IgM (μ) and other ELISA reagents were purchased from AbD Serotec. Human blood for lipid extraction was provided by local Guangzhou Blood Center. All other AR or higher grade chemicals were purchased from local chemical suppliers.

### 2.3. Sterile measures

The water used in all experiments was double distilled. All solutions or samples were sterilized by autoclaving or filtrating through 0.22 µm PVDF Syringe Filters. Solution or sample subpackage and mixing were performed in a laminar flow cabinet.

### 2.4. Extraction of erythrocyte lipid

Human or rat blood was diluted with normal saline and centrifuged at 4000rpm for 3 minutes to collect erythrocytes. The collected cells were washed by normal saline through 2-3 cycles of resuspension and centrifugation. The resuspension of washed erythrocytes was slowly dropped into a 10-fold volume of 0.2% acetic solution. The mixture was held at 4°C for hours to completely lyse erythrocytes and then centrifuged at 4000rpm for 10 minutes. The pellet was washed by 0.2% acetic solution through 2-3 cycles of resuspension and centrifugation to get white erythrocyte ghosts. A 20-fold volume of absolute ethanol was added into the erythrocyte ghosts. The mixture was shaken at intervals and held at 50°C in water bath for hours. Finally, erythrocyte lipid was obtained by centrifuging the mixture, recovering and drying the supernatant. Human erythrocyte lipid, rat erythrocyte lipid, and hemolytic rat erythrocyte lipid are, respectively, abbreviated as HEcL, REcL and LREcL. It was noted that the erythrocyte ghosts from severe hemolytic rat blood was slight brown and so LREcL showed deeper color than HEcL or REcL.

### 2.5. Preparation and quantification of hemozoin

Hemozoin was prepared following a previously published method with some modifications[48]. Briefly, 300mg of heme (hemin) was dissolved in 60ml of 0.1 M NaOH with stirring for 30 minutes. Glacial acetic acid was slowly dropped in to adjust the pH to about 4. Stirring was stopped and the mixture was heated at 70°C for 18 hours. After cooling, the separated solid was washed three times by 0.1M NaHCO_3_ for 3 hours and other three times by alternation of methanol and ddH_2_O. Finally, the solid hemozoin was dried in a drying oven overnight at 70°C. The quantification of hemozoin was conducted by redissolving it in 0.1M NaOH and by colorimetric method at 400nm with purity heme as the standard.

### 2.6. Formulations for immunizations

Phosphate Buffered Saline (PBS 1×) at pH7.4 was the basal solvent for all experiments and blank control. Heme solution was prepared by dissolving hemin in 0.1M NaOH to the concentration of 1mM for storage and diluted by PBS to a designed concentration for use. BSA (test antigen) solution was prepared by dissolving it in PBS to the concentration of 10mg/ml for storage and diluted by PBS to a designed concentration for use or negative control. BSA+CFA denotes the mixture of BSA solution and CFA and used for positive control. BSA+Heme denotes the mixture of BSA solution and heme solution. BSA+HEcL, BSA+REcL, BSA+LREcL, BSA+Heme+PC, BSA+Heme+REcL and BSA+Hz (hemozoin) +PC were all the form of liposome suspension and prepared briefly as follows. Lipid (0.1g) was completely dissolved in a 5ml mixture of ethanol and ether (1:1) in a 50 ml round bottom flask. In a fume hood, the flask was rotated slowly by hands while a nitrogen stream blew on the inner wall until a thin lipid film was formed and the solvent completely evaporated. BSA solution (2ml, 0.5mg/ml) with or without heme or hemozoin (in designed concentrations) was added into the flask. The flask was shaken to hydrate the lipid film to form multilamellar liposome suspension. Then the multilamellar liposome suspension was transferred into a 10ml conical flask to make small unilamellar liposome suspension by bath sonication. In most cases, the heme-contained formulations were liposome suspension because of the two reasons. First, the water solubility of heme is poor at the pH of 7.4. The application of lipid can make the formulations more stable. Second, the positive control CFA contains oil components. Meanwhile, hemolytic rat erythrocyte lipid (LREcL) is another positive control.

### 2.7. Immunizations and sera preparation

Male and female SD rats or BALB/c mice were obtained one week before immunizations and fed in a SPF environment. Rats or mice were grouped randomly and the number of each group was more than eight in case of accidental death by operations. Immunizations were conducted by intraperitoneal injection of 100µl formulation containing 50µg BSA for rats (42 days old) or 30µl formulation containing 15µg BSA for mice (42 days old). Blood was collected by cardiac or jugular venous puncture on the planned dates after immunizations. After blood clotting, the clear amber sera were collected by centrifugation at 4000rpm for 10 minutes and kept in 4°C to be tested.

### 2.8. Measurements of anti-BSA IgG and IgM

The levels of anti-BSA IgG and IgM were measured by indirect ELISA according to general guideline. Briefly, 96 well plates were coated with 10 µg/ml BSA overnight at 4C°. The coated plates were washed 3 times by wash buffer (PBS containing 0.05% Tween-20, 0.1ml/well). The plates were blocked with gelatin in PBS and incubated on a shaker for 2h at room temperature and then washed 3 times by wash buffer. Subsequently, a series of variously diluted rat or mouse sera were added as the first antibodies to the plates and incubated on a shaker for 1h at room temperature. After 3 times washing, HRP labeled ant-rat IgG (H/L), ant-rat IgM (μ), ant-mouse IgG (H/L) or anti-mouse IgM (μ) (diluted as the reagent manual) was added as the secondary antibody to the plates and incubated on shaker for 1h at room temperature. The plates were washed 3 times and TMB solution was added to conduct color development. After the reaction was stopped with 0.5 M H_2_SO_4_, the optical density was measured at 450nm (OD_450_) using an auto-plate reader. Each plate was read twice. The antibody standard control was the part of the protocol of the ELISA kit. The anti-BSA IgG or IgM level of each rat is indicated by the ELISA OD_450_ of the 100-fold diluted serum which lies in the range of nearly linear relationship between OD_450_ and dilution factor.

### 2.9. Spectroscopic measurements of BSA+Heme+PC

The preparation of BSA+Heme+PC was the same as in “*formulations for immunization*” except that the heme concentration of 40µM was much higher than those used in immunizations. If the concentration of 40µM cannot cause the conversion of heme into hemozoin, it is more impossible that those in immunizations result in the occurrence of hemozoin. To simultaneously measure the samples (liposome suspensions) that were held at 37°C for different days to make the errors as small as possible, the second sample was prepared 7 days post the first, and so on. On the thirty-fifth day, the seventh sample was done and all the samples were treated to disassemble liposomes to make the solutions clear by adding 2% SDS and shaking. The treated samples were centrifuged at 10000rpm for 10 minutes to check whether there were pellets (hemozoin crystals) in the tube bottoms. The supernatants were transferred for UV-VIS scanning from 300nm to 900nm.

## 3. Results

### 3.1. Lipid from hemolytic rat blood raises the level of anti-BSA antibodies in rats

Finding that there is IgM in Sprague Dawley (SD) rat sera against the antigens of ABO blood group (data not shown), we thought that human erythrocyte lipid (HEcL) with A or B antigen might have an adjuvant effect through the mediation of Fc receptors.

Investigation was performed to test whether HEcL can promote the production of anti-BSA antibodies in rats. In the experiments, the positive control was Complete Freund’s adjuvant (CFA), the negative control was rat erythrocyte lipid (REcL), the formulations were BSA-contained liposome suspension, and the administration was intraperitoneal injection. HEcL or REcL was prepared as the routine method: red cell ghosts were first made from fresh blood to separate cell membrane from hemoglobin and then the lipid was extracted from the ghosts. However, due to the mistake of substituting buffer for anticoagulant in drawing rat blood, the pooled rat blood was severe hemolytic and a lot of heme interfused in the cell membrane. So hemolytic rat erythrocyte lipid (LREcL) showed deeper color (slight brown) than HEcL. To our astonishment, the levels of anti-BSA IgG and IgM in the sera of the negative control group (immunized by BSA+LREcL liposome suspension) were approximated to those of the positive control group (immunized by BSA+CFA emulsion) on the 16^th^ day after immunizations (Fig. 1A-B). After repeating the experiments three times, including the extraction of LREcL on purpose, as well as studying related literatures, we deduced as follows. First, the results were not caused by LPS contamination or others because HEcL did not increase the level of anti-BSA IgG and IgM under the same experimental condition (Fig. 1A-B). Second, free heme may be the amazing actor because it is a DAMP to arouse innate immunity [36, 37] and innate immunity modulates adaptive immunity [46, 47]. Finally, the heme concentration that caused the results should be very low. It should be noted that concentration instead of amount/body weight is used for the dose of heme or BSA through this paper, because the immune response to intraperitoneal injection is firstly a local effect but not a whole body effect like to intravenous injection (the drug will be immediately throughout the body).

**Figure 1.**
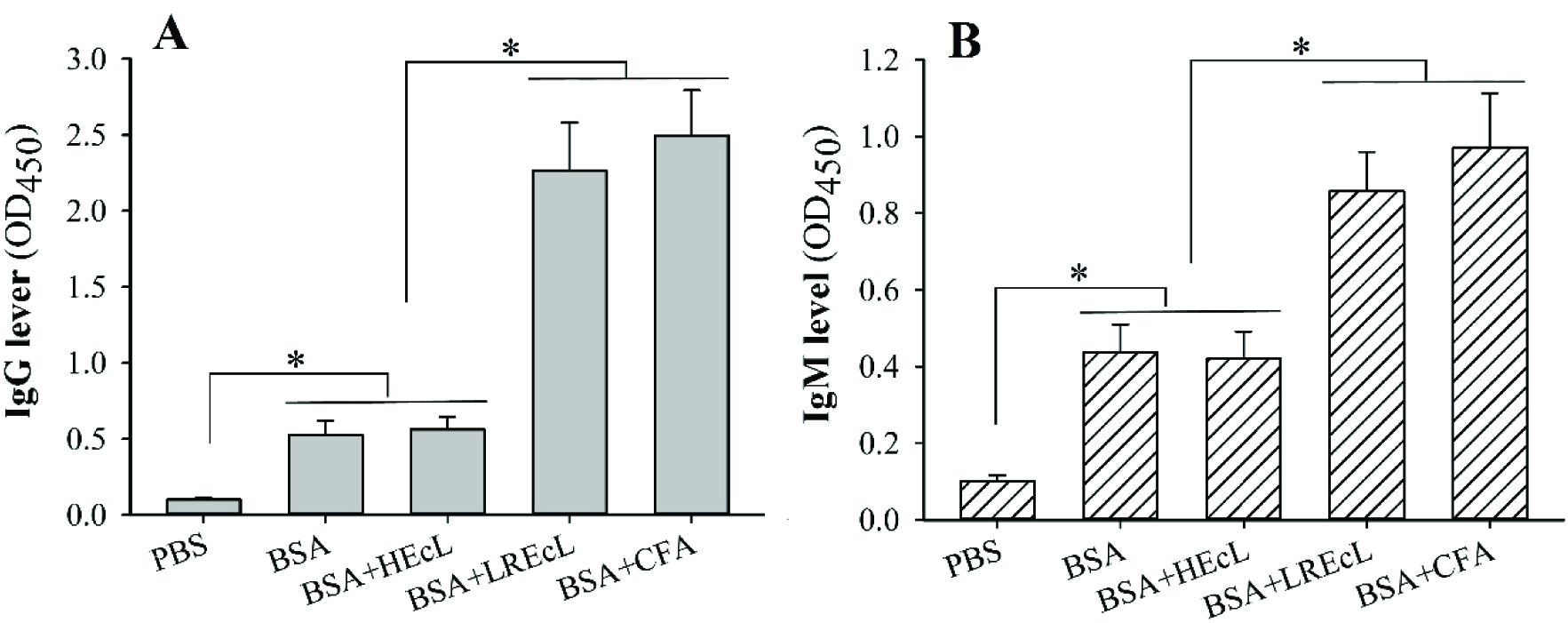
LREcL enhances anti-BSA antibody production in rats. Levels of anti-BSA IgG and IgM on the 16^th^ day after immunizations are indicated by the OD_450_ values of 100-fold diluted sera which are linearly related with the dilution factor. Immunizations were ip injection of 100 µl of formulations containing 50 µg BSA. Difference between each group (eight rats) is analyzed by t-test with two-tailed P-value. *, p < 0.001. BSA, bovine serum albumin. CFA, Complete Freund’s adjuvant. PBS, phosphate buffered saline. HEcL, human erythrocyte lipid. LREcL, hemolytic rat erythrocyte lipid. This figure was drawn from the data from one of the three repeated experiments.

### 3.2. Heme in several μM raises the level of anti-BSA antibodies in rats

To confirm whether heme is the enhancer in LREcL for the production of anti-BSA IgG and IgM in rats, commercial heme with the purity of HPLC grade (> 98%) was used. Limulus amebocyte lysate (LAL) test showed that the LPS activity was less than 0.01 EU/ml in 250 µM heme solution. In the experiments, heme-contained formulations were liposome suspension in most cases because of the two reasons. First, the problem stemmed from the LREcL which became a positive control in the subsequent experiments. Meanwhile, CFA, the positive control, also contains oil components. Second, the water solubility of heme is poor. The application of lipid (PC) can make the formulations more stable. The heme concentrations in all heme-contained formulations were adjusted by colorimetry to approximate to that in LREcL (about 2.3 µM). On the fifteenth day after immunizations, compared to the formulations without heme, all those with heme significantly raised the levels of anti-BSA IgG and IgM in rat sera (Fig. 2A-B).

**Figure 2.**
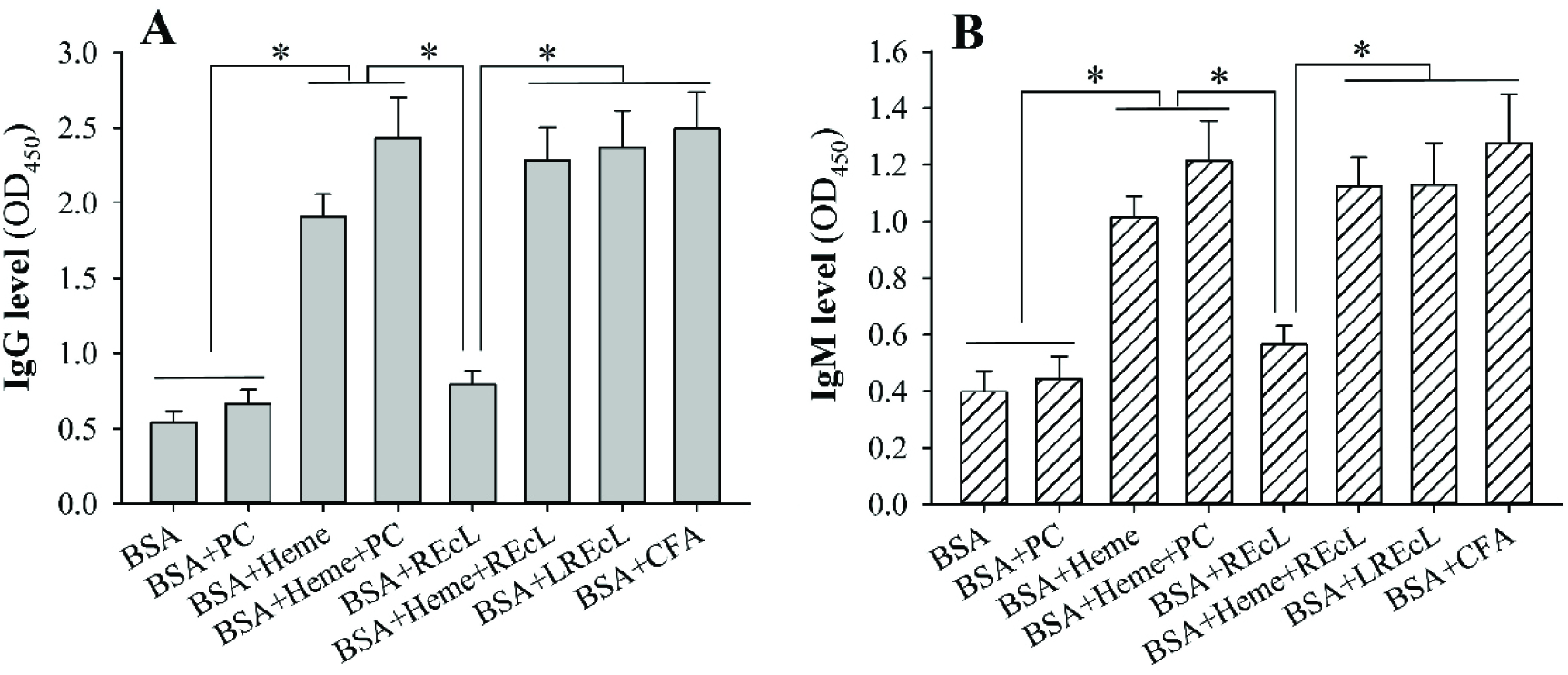
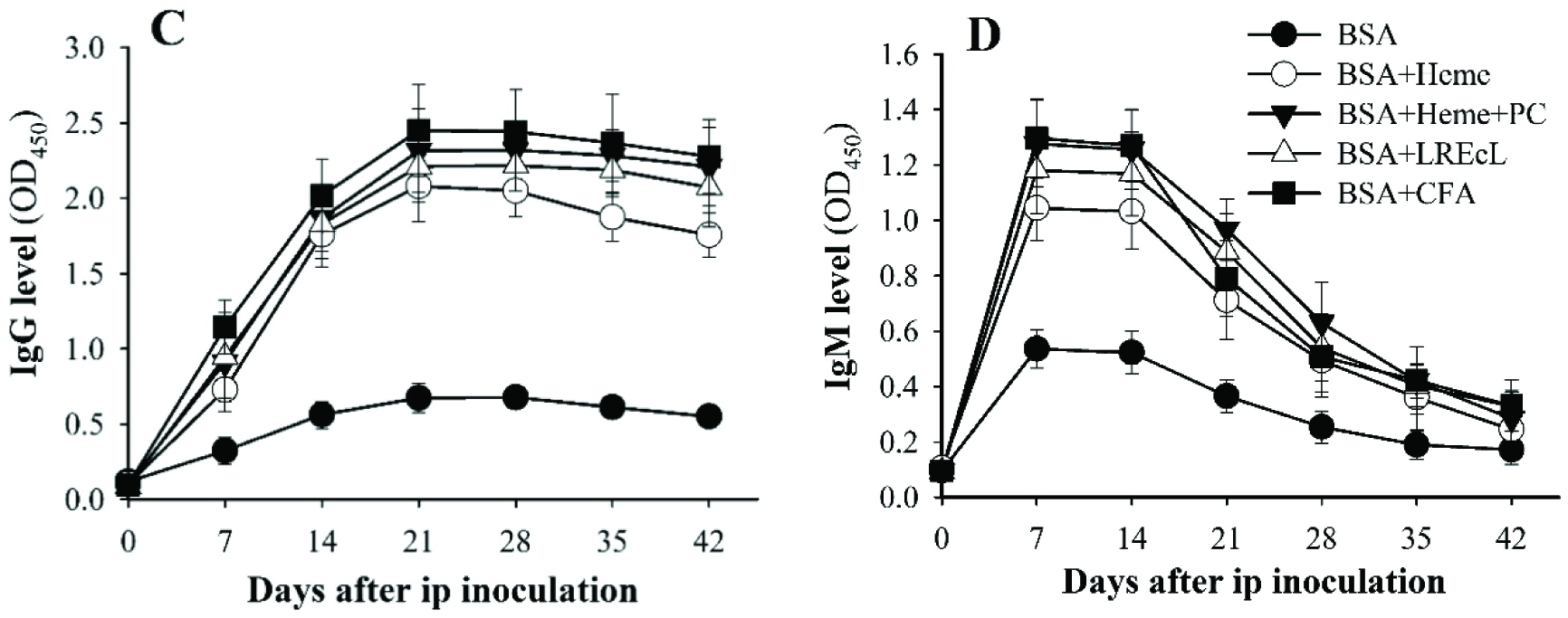
Heme in several μM enhances anti-BSA antibody production in rats. (A-B) Levels of anti-BSA IgG and IgM on the 15^th^ day after immunizations. (C-D) Time kinetics of the production of anti-BSA IgG and IgM. Anti-BSA levels are indicated by the OD_450_ values of 100-fold diluted sera which are linearly related with the dilution factor. Immunizations were ip injection of 100 µl of formulations containing 50 µg BSA. The heme concentrations in all heme-contained formulations were about 2.3 µM adjusted by colorimetric comparison with LREcL sample. Difference between each group (eight rats) is analyzed by t-test with two-tailed P-value. *, p < 0.001. BSA, bovine serum albumin. CFA, Complete Freund’s adjuvant. PBS, phosphate buffered saline. PC, phosphatidylcholine. REcL, rat erythrocyte lipid. LREcL, hemolytic rat erythrocyte lipid. This figure was drawn from the data from one of the two repeated experiments.

Furthermore, except heme alone, the promotive ability of heme+PC or heme+REcL was approximated to that of CFA or LREcL. In other independent experiment, the time kinetics of the production of anti-BSA IgG and IgM in the presence of heme was parallel to that in the presence of CFA or LREcL (Fig. 2C-D). The results demonstrate that heme, at least in very low concentrations and by intraperitoneal injection, can increase the production of specific antibodies in rats.

### 3.3. Heme modulates the production of anti-BSA antibodies in mice by the mode of double concentration-thresholds

The above results encouraged us to study the dose effect of heme (plus PC) on the production of ant-BSA IgG and IgM by mouse experiments. Firstly, it was performed to find out the upper concentration limit at which heme stops its promotion. However, to our astonishment again, the levels of anti-BSA IgG and IgM in the group of high heme concentrations were even below those in the group of BSA alone. We realized that heme at high concentrations exerts a suppressive effect on the production of anti-BSA antibodies. To find the concentration intervals where heme may exert different effects on the production of anti-BSA antibodies, a series of heme concentrations were designed from 0.1 to 15µM. Compared to BSA+PC, the formulations at the heme concentrations below 1µM (the first threshold) did not raise the levels of anti-BSA IgG and IgM. However, when the heme concentrations were more than 5µM (the second threshold), the levels of anti-BSA IgG and IgM fell down below the levels of the group of BSA alone. Only between 1 µM and 5 µM, a concentration interval, the levels of anti-BSA IgG and IgM were significantly increased (Fig. 3A-B). The results demonstrate that heme can regulate animal adaptive immunity (at least humoral immunity) by the mode of double concentration-thresholds. If heme concentrations are below the first threshold, the effect of heme on adaptive immunity is zero; if between the first and second thresholds, the effect is promotive; if over the second threshold, the effect is inhibitive.

**Figure 3.**
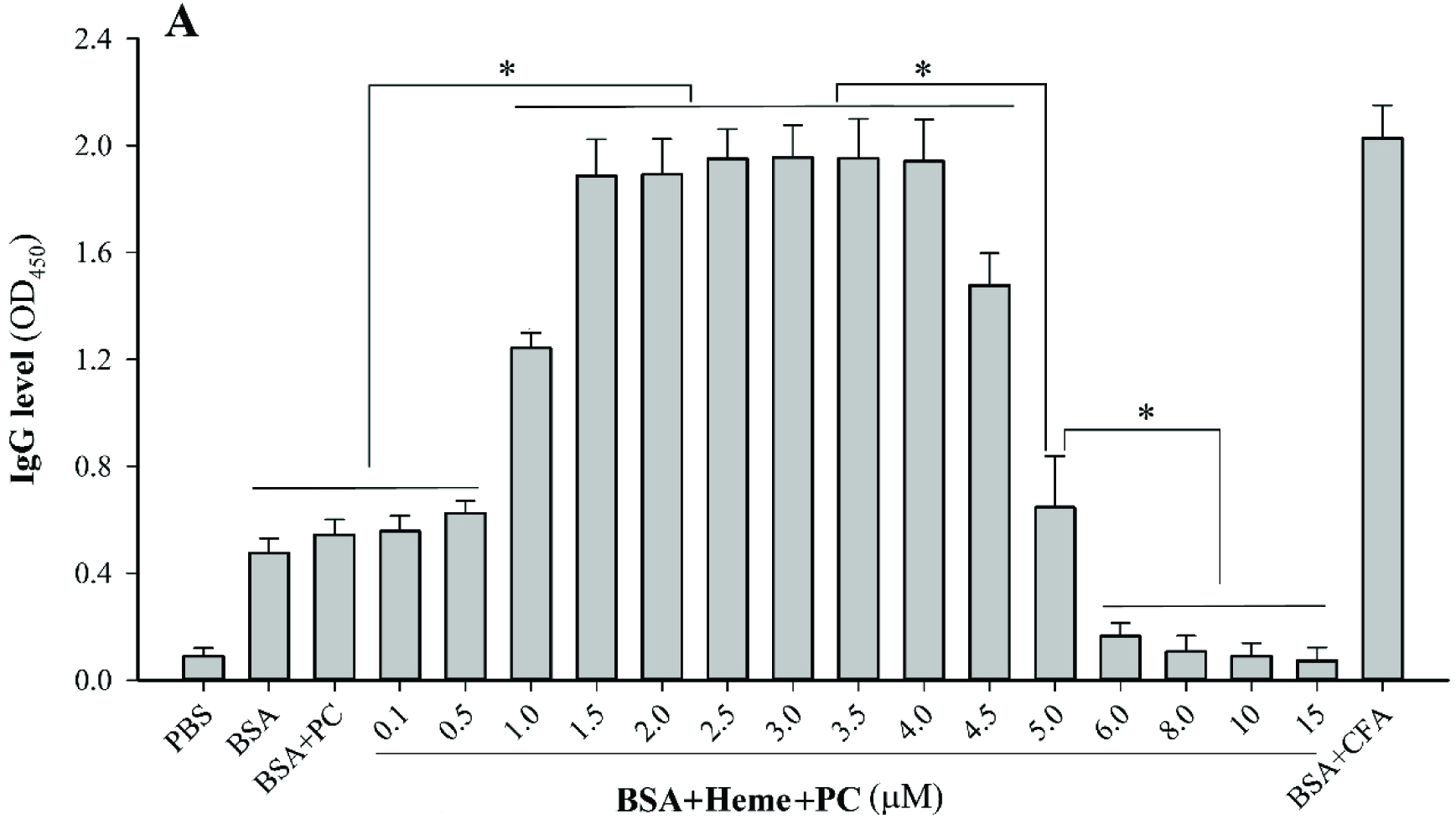
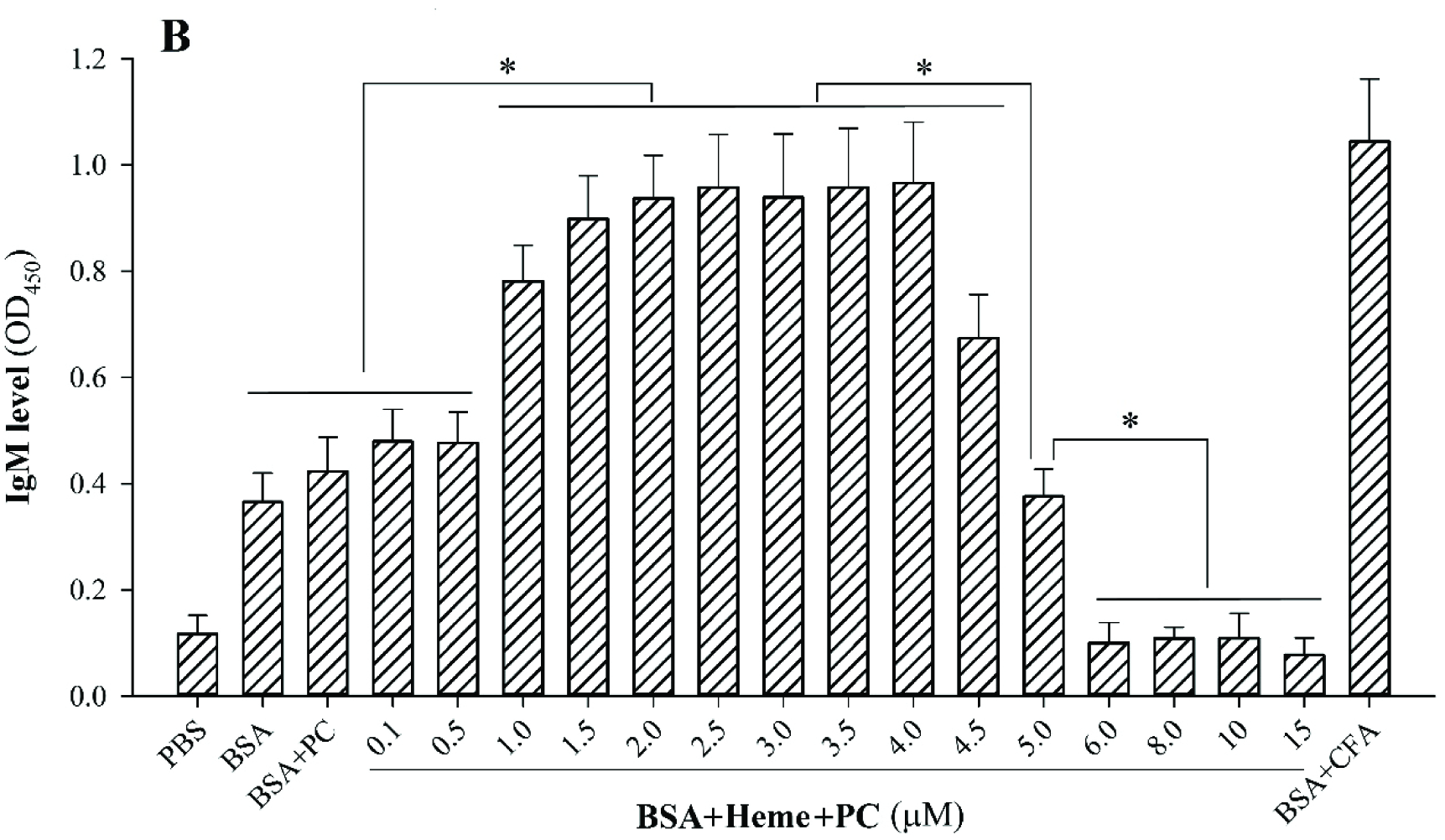
Heme concentration-dependently modulates anti-BSA antibody production. Levels of anti-BSA IgG and IgM in mice on the 15^th^ day after immunizations are indicated by the OD_450_ values of 100-fold diluted sera which are linearly related with the dilution factor. Immunizations were ip injection of 30 µl of formulations containing 15µg BSA. Difference between each group (eight mice) is analyzed by t-test with two-tailed P-value. *, p < 0.001. BSA, bovine serum albumin. CFA, Complete Freund’s adjuvant. PBS, phosphate buffered saline. PC, phosphatidylcholine. This figure was drawn from the data from one of the two repeated experiments.

### 3.4. Enhanced production of anti-BSA antibodies is not from heme-derived hemozoin during immunizations

Hemozoin is heme-derived insoluble crystals, causes innate immunity/inflammation, and has been applied as an adjuvant[48, 49]. So the conversion of heme into hemozoin must be excluded during immunizations. As far as is known, the pH and heme concentration for hemozoin occurrence are less than 5 and more than 100µM, respectively[48]. Although the pH (7.4) and heme concentrations (0.1-15µM) in our experiments were theoretically impossible for hemozoin occurrence, *in vivo* and *in vitro* experiments were performed to verify whether heme transformed into hemozoin during the immunizations. In *in vivo* experiments, the concentrations of hemozoin (insoluble nanoparticles) used in the immunization formulations were equivalent to the concentrations of heme from 50 to 200µM. If the transformation of heme into hemozoin was true during the immunizations, as heme in the concentrations over 6µM did (Fig. 3A-B), hemozoin in the concentrations over 50µM should suppress the production of anti-BSA antibodies. However, hemozoin in the concentrations over 50µM enhanced the production of anti-BSA IgG and IgM (Fig. 4A-B). In *in vitro* experiments, the formulations with the heme concentration of 40µM were held at 37 °C for different days. If heme in the concentrations from 0.1 to 15µM can transform into hemozoin during immunizations, heme in the concentration of 40µM should more preferentially to form hemozoin. However, regardless of how many days the samples were hold for, after centrifugation, there were no pellets (hemozoin nanoparticles) in the tube bottoms. Furthermore, the spectra of the supernatants were almost the same, which meant no hemozoin occurrence (Fig. 4C). The results of the *in vivo* and *in vitro* experiments demonstrate that heme cannot transform into hemozoin during immunizations. That is to say, the production of anti-BSA IgG and IgM in our experiments was enhanced by heme but not by hemozoin.

**Figure 4.**
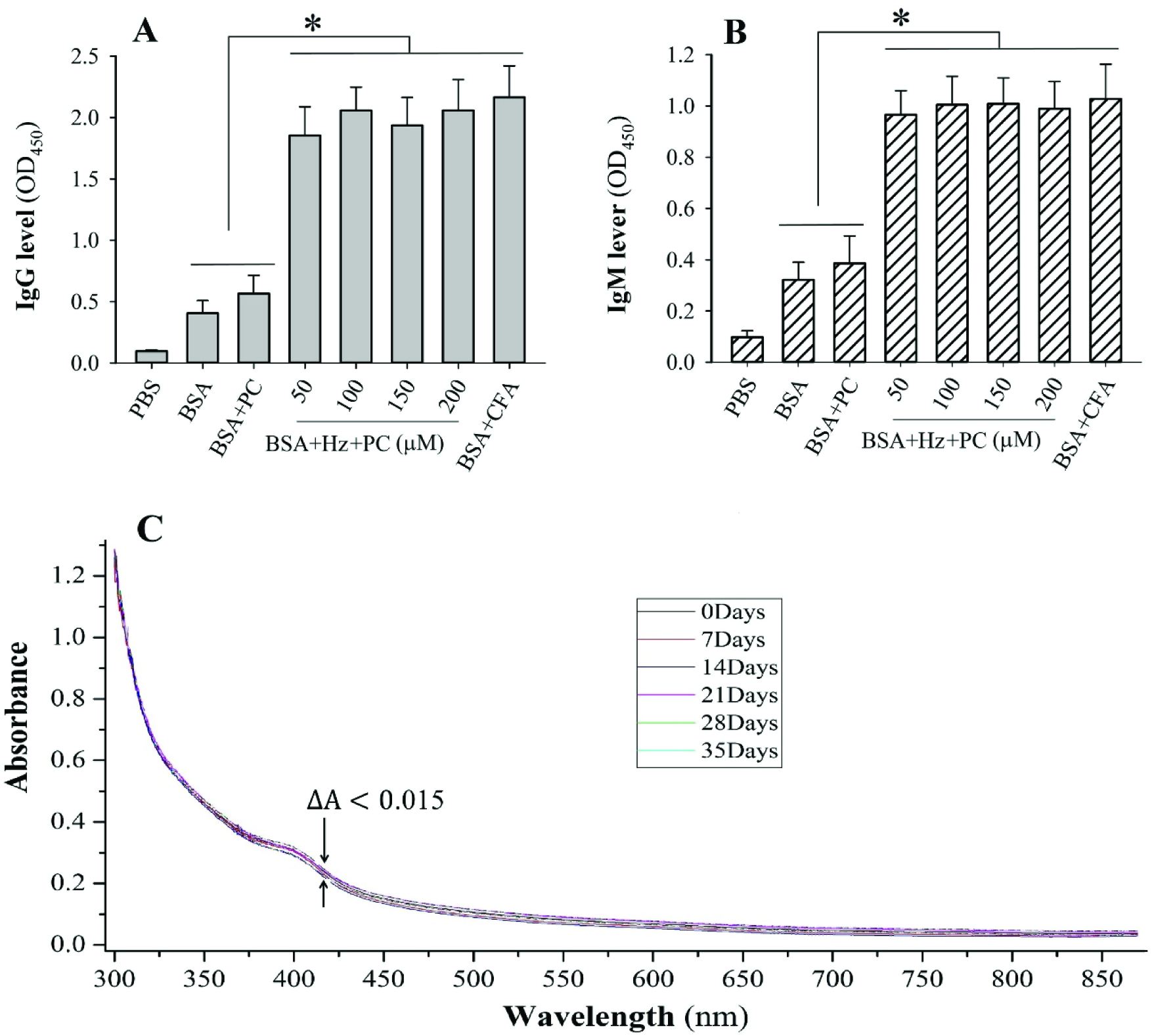
Enhanced anti-BSA antibody production are not caused by the conversion of heme into hemozoin (Hz) during immunizations. (A-B) Levels of anti-BSA IgG and IgM on the 15^th^ day after immunizations. Anti-BSA Levels are indicated by the OD_450_ values of 100-fold diluted sera which are linearly related with the dilution factor. Immunizations were ip injection of 30µl of formulations containing 15µg BSA. Being insoluble, hemozoin concentration is indicated by the equivalent of heme. Difference between each group (eight mice) is analyzed by t-test with two-tailed P-value. *, p < 0.001. BSA, bovine serum albumin. CFA, Complete Freund’s adjuvant. PBS, phosphate buffered saline. PC, phosphatidylcholine. (C) Spectra of heme in BSA+heme+PC samples that were held at 37°C for different days. The heme concentration was 40µM.

## 4. Discussion

### 4.1. Questions about LPS contamination and binding of heme to BSA

LPS is a TLR4 activator and usually used as an adjuvant ingredient[50]. Free heme can synergize with low concentrations of LPS [26]. So there are two questionable points. The enhancer for the production of anti-BSA antibodies may be LPS contamination but not heme. Even if the LPS contamination is too less to show its effect, the enhancer may be the synergy of heme with LPS instead of heme alone. The two possibilities can be excluded by the following facts. First, besides heme being high purity (> 98%) and qualified with LAL test, all samples were prepared under sterile conditions to prevent them from any contaminations. Second, if the enhancer is LPS contamination, under the same experimental condition, all the samples should be at equal chance contaminated by LPS and show the same promotive effect. However, the enhanced production of anti-BSA antibodies always happened only in some certain samples (Fig. 1-4). In the third place, If the enhancer is the synergy of heme with low concentrations of LPS contamination, the higher heme concentrations should at least show the same effect as the lower heme concentrations do. However, unlike lower heme concentrations (1-5µM) to promote the production of anti-BSA antibodies, the higher heme concentrations (>5µM) suppressed the production of anti-BSA antibodies (Fig. 3).

The unique chemical structure makes heme easily bind with proteins through diverse ways, including coordination of the central iron ion with N, S or O atom on the side chain of His, 300 Lys, Cys, Met or Tyr; hydrophobic interaction of porphyrin ring, methyl and vinyl groups with non-polar residues; electrostatic interaction of propionate groups with positive charged residues and covalent bond of vinyl groups with Cys residues. So a question arises. Did the binding of heme to BSA but not free heme enhance the antigenicity of BSA? This assumption conflicts with the following facts. First, it was reported that free heme but not protein-bound heme can stimulate immune system[51]. Second, heme is easy to dissolve in lipid (or insert into membrane), which will impair the binding of heme to BSA. So, if the assumption is true, the antibody levels induced by BSA+heme should not markedly lower than those induced by BSA+heme+lipid under the same concentrations of heme and BSA. But the fact was opposite to the assumption (Fig. 2A-B & Fig. 3). Finally, higher heme concentrations will promote the binding of heme to BSA. So, if the assumption is true, the higher heme concentrations should at least show the same effect as the lower heme concentrations do. However, unlike lower heme concentrations (1-5µM) to promote the production of anti-BSA antibodies, the higher heme concentrations (>5µM) suppressed the production of anti-BSA antibodies (Fig. 3).

### 4.2. Double concentration-thresholds mode of heme modulating immune system

Our experimental results demonstrate that heme can modulate adaptive immune response by the mode of double concentration-thresholds. In detail, when the concentrations of used heme are below the first threshold, heme shows no effect on adaptive immune response; when between the first and second thresholds, heme shows promotive effect; when over the second threshold, heme shows inhibitive effect. This mode suggests that physiological release of heme may be one of the intrinsic mechanism for animals to maintain an adequate immunity while pathological bleeding may result in animal immunosuppression. It is note that this regulation should only run outside the circulatory system because there is a powerful buffer system (haptoglobin, hemopexin and other serum proteins) against free heme in the circulatory system[2].

Why can heme work in this way? Although the concrete molecular mechanism is to be explored, according to previous related reports, we present the following hypothesis (Fig. 5). As described in introduction, immune system is antagonistically affected by two heme-led actions: heme action and HO action. Through activating TLR4 dependent[21-23] and ROS dependent signaling pathways[24-26], free heme acts to potentially enhance adaptive immunity[46, 47]. Meanwhile, by degrading heme, HO (including inducible HO-1 and constitutive HO-2) acts to set up an immunosuppressive mechanism triggered by the catabolites CO, biliverdin and bilirubin[41-43]. Heme action and HO action are not independent but interact with each other. First, heme is their common pilot. Second, heme induce the expression of HO-1 to reinforce HO action[44]. Finally, HO depletes heme by degrading it and the degradation products inhibit TLR4 signaling pathway as well as scavenge ROS[52].

**Figure 5.**
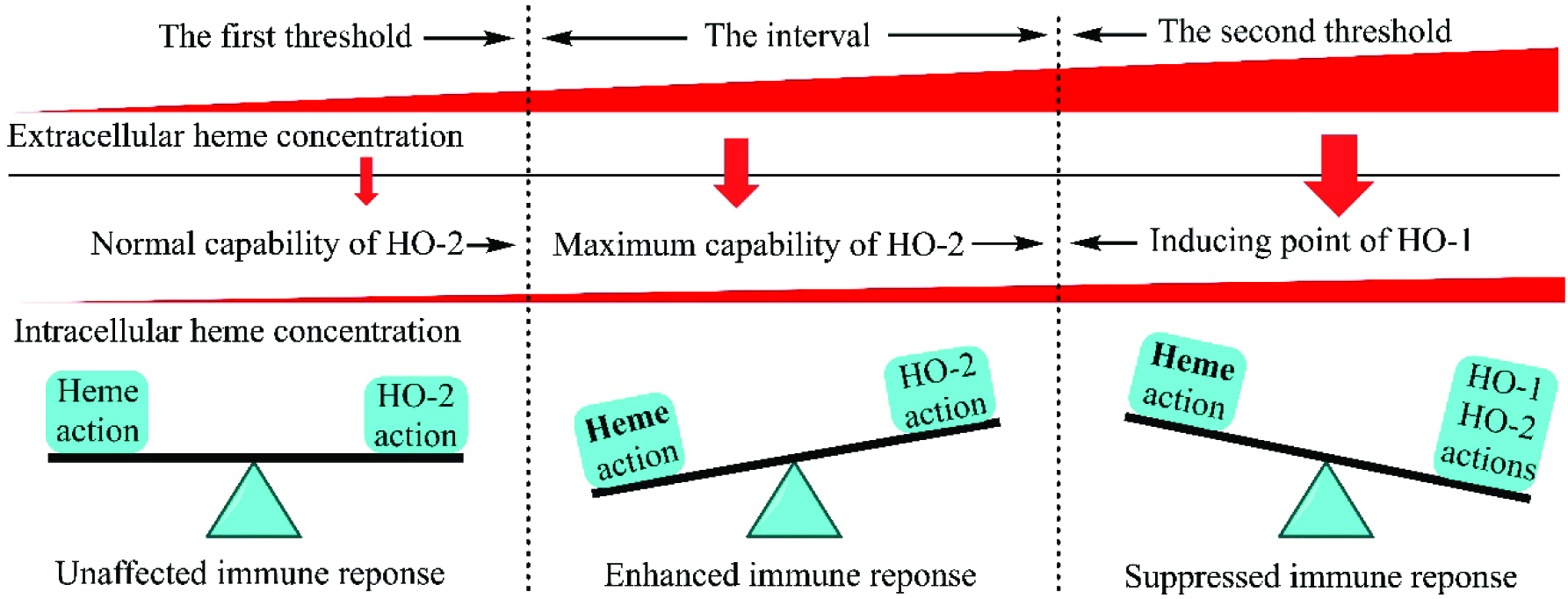
Possible mechanism of the mode of double concentration-thresholds.

The competition and crosstalk between heme action and HO action defines the mode of double concentration-thresholds (Fig. 5). When the concentrations of free heme outside immunocytes are below the first threshold, the concentrations of free heme inside immunocytes will not be over the normal capability of HO-2. Under this condition, heme action and HO-2 action are balanced and so adaptive immunity is unaffected. When the extracellular concentrations of free heme are between the first and second thresholds, the intracellular concentration of free heme will be more than the normal capability of HO-2 but not be able to induce the expression of HO-1. In this case, heme action exceeds HO-2 action and therefore adaptive immunity is promoted. When the extracellular concentrations of free heme are more than the second threshold, the intracellular concentrations of free heme will be able to induce the expression of HO-1. Certainly, the sum of HO-1 action and HO-2 action surpasses heme action and thus adaptive immunity is suppressed.

### 4.3. Immunological significance of this finding

Animal immune system can recognize both the exogenous and endogenous danger signals, including pathogen-associated molecular patterns (PAMPs) and damage-associated molecular patterns (DAMPs)[53]. PAMPs and DAMPs activate pattern recognition receptors dependent signaling pathways to provoke innate immunity and to further regulate adaptive immunity[46, 47]. Many PAMPs or their analogues have been applied in the design of immunoadjuvants and vaccines [50]. However, there have been so far no reports about the effect of a DAMP on adaptive immune response to an antigen. Free heme is an endogenous danger signal, a DAMP to provoke innate immunity[36, 37]. As far as we know, it should be the first report that free heme regulated the production of anti-BSA antibodies in rats and mice by the mode of double concentration-thresholds. This finding suggests that free heme can concentration-dependently modulate animal adaptive immune response to an antigen and DAMPs such as heme, like PAMPs, can be used as a basic ingredient of adjuvants.

## Acknowledgements

This work was supported by the National Natural Science Foundation of China (Grant No. 366 30371326), and the Natural Science Foundation of Guangdong, China (Grant No. 367 8151027501000007).

